# Numerical study of the formation and stability of a pair of particles of different sizes in inertial microfluidics

**DOI:** 10.1101/2022.12.14.520448

**Authors:** Krishnaveni Thota, Benjamin Owen, Timm Krüger

## Abstract

The formation of pairs and trains of particles in inertial microfluidics is an important consideration for device design and applications, such as particle focussing and separation. We study the formation and stability of linear and staggered pairs of nearly rigid spherical particles of different sizes in a pressure-driven flow through a straight duct with rectangular cross-section under mild inertia. An in-house lattice-Boltzmann-immersed-boundary-finite-element code is used for the simulations. We find that the stability and properties of pairs of heterogeneous particles strongly depends on the particle sizes and their size ratio, while the formation of the pairs is also determined by the initial lateral position and the axial order of the particles. Our findings imply that perturbations of particle trajectories caused by other particles, as they are expected to happen even in dilute suspensions, can be important for the formation of stable pairs in inertial microfluidics.

## I. INTRODUCTION

Separation and sorting of micron-sized particles and cells has importance in disease diagnostics^1^, therapeutics^2^, and cell analysis^3^. Among the available separation methods, inertial microfluidics (IMF) has become attractive due to its high throughput, low cost, and label-free manipulation of the particles^4^. IMF is a relatively new field that emerged in the late 2000s^5,6^. While inertia is often negligible in traditional microfluidic devices due to the small length scales and flow rates involved, the channel Reynolds number in IMF is of the order 10-100 due to the relatively high velocity of the fluid. In this range of Reynolds number, inertial effects can be exploited to manipulate particles through focussing and separation^7–10^.

Particles in IMF experience shear gradient and wall repulsion forces^11–14^. The shear gradient lift force results from the interaction of the finite size of the particle with the gradient of the flow velocity across the channel. This force usually pushes the particle away from the channel centre towards a wall. The wall repulsion force is caused by an increased pressure between the particle and the wall. The resulting net force felt by the particle is commonly known as inertial lift force. As a consequence, particles usually undergo lateral migration towards one of the existing lateral equilibrium positions. This particle focussing phenomenon was first observed by Segré and Silberberg^15^ in a pressure-driven flow through a cylindrical pipe.

In addition to the focusing of a single particle in IMF, multiple particles tend to form axially ordered trains with regular inter-particle spacing^16,17^. The formation of trains can be exploited in applications such as cell encapsulation^18^ and flow cytometry^19^. Since it has been identified that the formation of particle pairs precedes the emergence of trains^20^, understanding the formation of particle pairs is crucial. Lee *et al*.^21^ first identified the self-assembly of particle pairs and identified reverse streamlines are crucial elements. Particle pairs can be classified into staggered pairs (particles located on opposite sides of the channel) and linear pairs (particles placed on the same side of the channel)^20,21^. During pair formation, the axial distance between both particles performs a damped oscillation before converging to an equilibrium value^16,22^. It has been reported that linear particle pairs do not form when both particles are of the same size^20^.

Patel and Stark^23^ investigated the effect of particle softness and shape for mono- and bi-disperse particle pairs and found that the presence of the second particle can change the stability of the single-particle equilibrium positions. Li *et al*.^24^ investigated the formation of a heterogeneous pair of particles consisting of a rigid and a soft particle and demonstrated the pair formation after a number of passing interactions in a simulation with periodic boundary conditions. Gao *et al*.^25^ experimentally found that a small difference in particle size improves the particle focussing performance. Chen *et al*.^26^ performed 2D simulations of the pair formation of bi-dispersed particles of different sizes in a linear arrangement; they found that pair formation is only possible when the larger particle is leading.

The formation of heterogeneous pairs can be desired (*e.g*., for the generation of Janus or compound particles) or detrimental *(e.g*., for the separation of different particles). Thus, it is important to better understand the conditions and mechanisms leading to the formation of heterogeneous pairs. In this paper, we use a lattice-Boltzmann-immersed-boundary-finite-element solver to numerically investigate the dynamics and formation of a pair of particles of different sizes through a straight rectangular channel by a pressure-driven flow at a moderate Reynolds number (Sec. II). We consider particle pairs in the staggered and linear arrangements (Sec. III). Our study comprises three parts. First, we investigate for which combinations of particle sizes stable pairs form when both particles are initially far away from each other and at their respective lateral equilibrium positions; we find that the formation of these ‘unperturbed pairs’ not only depends on the size ratio, but also on the absolute sizes of the particles. Interestingly, the known instability of linear pairs of identical particles disappears when there exists a small size heterogeneity. Second, we analyse the stability of already existing pairs, independently of their possible formation mechanism; we observe that more pairs are stable than those that form under unperturbed conditions. Third, we study how a perturbation of the initial position of the smaller particle, which might be caused by the presence of other particles, affects pair formation; we identify the existence of a ‘stability band’, a finite region of initial positions of the smaller particle that lead to stable pairs. Implications and future directions are discussed in Sec. IV.

## II. PHYSICAL MODEL AND NUMERICAL METHODS

We study, via computer simulations, the dynamics of a pair of nearly rigid spherical particles of different sizes in a flow through a straight microchannel at moderate Reynolds numbers. In the following, we briefly explain the physical model (Sec. II A), including the geometrical setup, and the numerical methods (Sec. II B).

### A. Physical model

We consider a Newtonian liquid with kinematic viscosity v and density *ρ* flowing through a rectangular duct with a width of 2*w* and a height of 2*h* with an aspect ratio *w/h* = 2 as shown in Fig. 1a. The liquid is governed by the incompressible Navier-Stokes equations. Flow is driven along the *x*-axis (axial direction). The *y*- and *z*-axes are denoted as lateral directions.

**FIG. 1:**
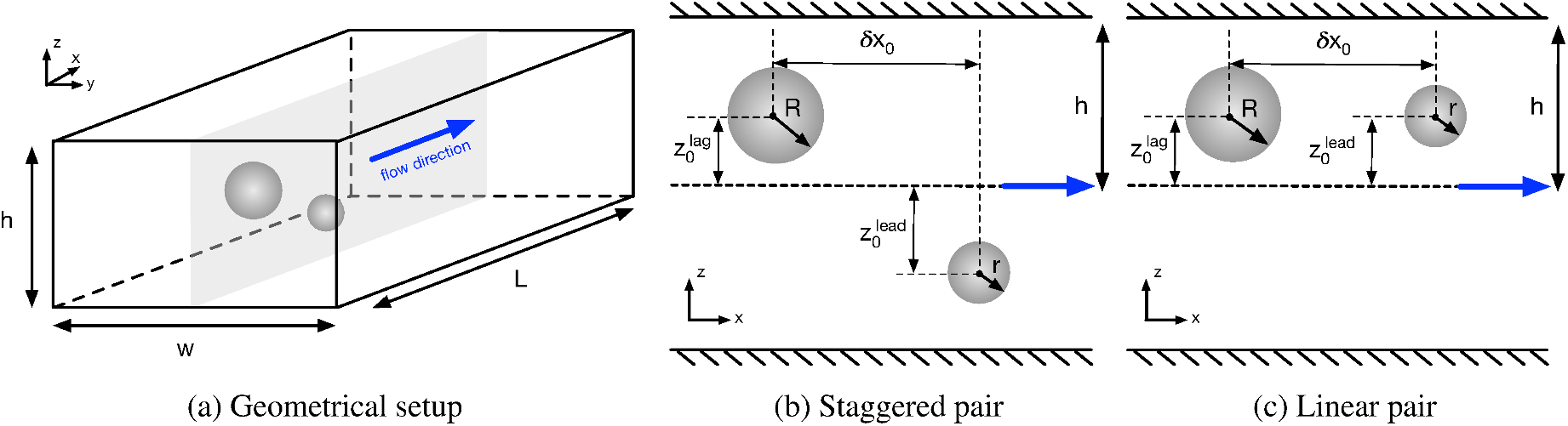
(a) Schematic of a pair of particles in a rectangular duct with height 2*h* and width 2*w*; the length of the periodic unit cell is *L*. The flow is along the *x*-axis (blue arrow). The particles are initially located on the *x*-*z*-mid-plane (*y* = 0, grey plane), (b) In the staggered arrangement, both particles are initially on different sides of the channel. (c) Particles in the linear arrangement are initially on the same side of the channel. The initial axial distance between the particles is *δx*_0_ ≪ *L*. The particle initially located downstream is called leading particle; the other particle is called lagging particle.

We consider two initially spherical and neutrally buoyant particles with radii *R* and *r* ≤ *R*. The size ratio of the two particles is *β* = *R/r* ≥ 1. Note that in some cases we consider the limit of homogeneous particles, *β* = 1. Both particles are modelled as capsules comprising a thin hyperelastic membrane and an interior liquid with the same properties as the suspending liquid. The capsule membranes are governed by the Skalak model^27^:

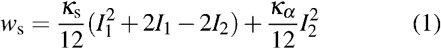

where *w_s_* is the areal energy density, *I*_1_ and *I*_2_ are the in-plane strain invariants^28^, and *κ*_s_ and *κ*_α_ are the elastic shear and area dilation moduli. In order to avoid membrane buckling, we include a membrane bending energy

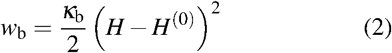

where *H* and *H*^(0)^ are the trace of the surface curvature tensor and the spontaneous curvature, respectively, and *κ*_b_, is the bending modulus.

Both particles are initially placed on the *x*-*z*-mid-plane (*y* = 0) between the side walls as shown in Fig. 1a. Particles initially located on this mid-plane will usually stay on the plane while moving along the *x*-axis and migrating along the *z*-axis^29^. We distinguish between the initially lagging and leading particles based on their initial positions on the *x*-axis (Fig. 1b and Fig. 1c). We consider two arrangements of particles in this work. In the first arrangement, the particles are placed on opposite sides of the channel centre (staggered arrangement, Fig. 1b). In the second arrangement, the particles are positioned on the same side of the channel (linear arrangement, Fig. 1c). In both arrangements, we distinguish between cases with the smaller particle being the leading or the lagging particle.

We apply periodic boundary conditions in the axial direction and the no-slip condition at the channel walls and on the surface of the particles. The channel length *L* is large enough so that particles do not interact with their periodic images.

The channel Reynolds number Re is defined as

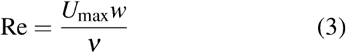

where *U*_max_ is the maximum velocity of the flow in the absence of particles. Following Schaaf and Stark^30^, the Laplace number La is used to characterise the particle softness:

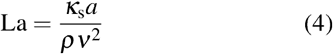

where *a* is the radius of the particle which is either *R* or *r*.

We use the channel half-height *h* as characteristic length to non-dimensionalise the particle radii and the travelled distance. Time is non-dimensionalised by the advection time of the larger particle:

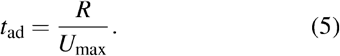

Other dimensionless groups are the confinement of the larger particle, *χ_R_* = *R/h*, the confinement of the smaller particle, *χ_r_* = *r/h*, the channel aspect ratio *α* = *w/h*, the reduced dilation modulus 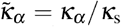, and the reduced bending modulus 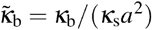.

### B. Numerical model

We use a fluid-structure interaction solver in which the lattice-Boltzmann (LB) method is used to solve the Navier-Stokes equation, the finite-element (FE) method for the particle dynamics, and the immersed-boundary (IB) method for the fluid-structure interaction. This IB-LB-FE solver has been previously employed in the study of deformable capsules in inertial microfluidics^31,32^· We only provide essential properties of the model here, while comprehensive details are available elsewhere^28^.

We use the D3Q19 lattice^33^ and the BGK collision operator^34^ with relaxation time τ for the LB method. The viscosity of the liquid and the relaxation time satisfy

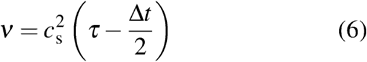

where *c_s_* is the lattice speed of sound and Δ*t* is the time step. For the D3Q19 lattice, 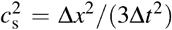 holds where Δ*x* is the lattice resolution. The flow is driven by a constant body force^35^. The no-slip boundary condition at the channel wall is realised by the standard half-way bounce-back condition^36^. This form of the LB method is widely used in the field of fluid dynamics, including in previous inertial microfluidics studies^30,37^.

A surface mesh consisting of flat triangular faces (or elements) defined by three nodes (or vertices) each is used to represent each particle. The particle mesh is generally deformed at a given time step. An explicit scheme is used to calculate the resulting hyperelastic forces acting on each vertex. The bending forces are calculated from the angles between normal vectors of pairs of neighbouring faces, and the shear and area dilation forces are calculated from the deformation gradient tensor of each face^38^.

We employ an IB method with a 3-point stencil^39^. The forces obtained from the FE scheme are spread from the Lagrangian particle mesh to the Eulerian fluid lattice where they act on the surrounding fluid nodes through the LB algorithm. The updated fluid velocity is then interpolated at the location of each mesh node. The positions of the mesh nodes are updated using the forward-Euler method, assuming a massless membrane which is appropriate for neutrally buoyant capsules. This treatment recovers the no-slip boundary condition at the surface of the capsules and the momentum exchange between the liquid and the particles.

Our IB-LB-FE solver has been tested for a single soft particle and the interaction between two soft particles in inertial flows in our previous work^32^. Unless otherwise stated, the following parameters are kept constant in this study: 2*w* = 160Δ*x*, 2*h* = 80Δ*x*, *L* = 560Δ*x*, Re = 10, La = 100, 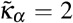, and 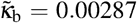. The number of surface mesh elements of the particles range from 500 for the smallest simulated particle to 7220 for the largest one. We focus on *R*, *r*, *δx*_0_, 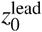 and 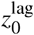 as free parameters. Note that, for La = 100, particles are close to the rigid limit^32^. Simulations are initialised by dropping the particles in the simulation box and then driving the flow, starting at *t* = 0.

## III. RESULTS AND DISCUSSIONS

We first explain the types of pair interaction observed in our study together with examples of their trajectories. We then investigate the effect of the size and initial lateral position of the particles on the formation and stability of particle pairs.

### A. Types of observed particle pairs

In a Poiseuille flow of a Newtonian liquid at mild inertia, a suspended rigid spherical particle generally migrates to a lateral equilibrium position between the channel centre and the wall^5^. Larger particles are usually focussed at a lateral position closer to the centreline than smaller particles^40,41^. Due to the curved velocity profile, particles focussed closer to the centreline generally move faster along the channel than those focussed nearer to a wall. Hence, to leading order, stable pairs are only expected to exist when both particles have the same axial speed at their respective equilibrium positions. However, the presence of a second particle nearby can modify the lift force experienced by the first particle — and vice versa^37^. Due to the hydrodynamic interaction between both particles, lateral equilibrium positions can change and the particles can form a stable pair under some circumstances, even if both particles have different individual lateral equilibrium positions.

Not all pairs exhibit the same characteristics, and pairs can be categorised according to the time evolution of their axial distance. We observe four different types of interactions between two particles: stable pairs, oscillatory stable pairs, unstable pairs, and periodic pairs. We define a stable pair as an arrangement where the axial distance between the particles, *δx*. converges to a constant value which is sufficiently small so that particles still interact hydrodynamically (typically *δx/R* < 6); we denote this equilibrium axial distance *δx*_eq_.

An example of a stable pair in the staggered arrangement is shown in Fig. 2a for particles with *χ_R_* = 0.35 and *χ_r_* = 0.3. The particles were initialised at their single-particle equilibrium positions with the larger particle lagging behind the smaller particle by an axial distance *δx*_0_/*R* = 10. The larger particle is closer to the centre and, therefore, moves with a higher axial speed. Initially, the larger particle approaches the leading smaller particle and attempts to overtake; the axial distance decreases. During this process, the lateral positions of both particles change in a way that the smaller particle gets closer to the centre than the larger particle. Consequently, the larger particle fails to overtake the smaller particle, and the smaller particle moves faster than the larger particle for some time. In the following, both particles tend to migrate back to their respective lateral equilibrium positions, thus the larger particle becomes faster and the axial distance decreases again. The process repeats for a few times with a decreasing amplitude, until both particles settle in a stable arrangement where both particles move with the same axial speed and the smaller particle is still leading.

**FIG. 2:**
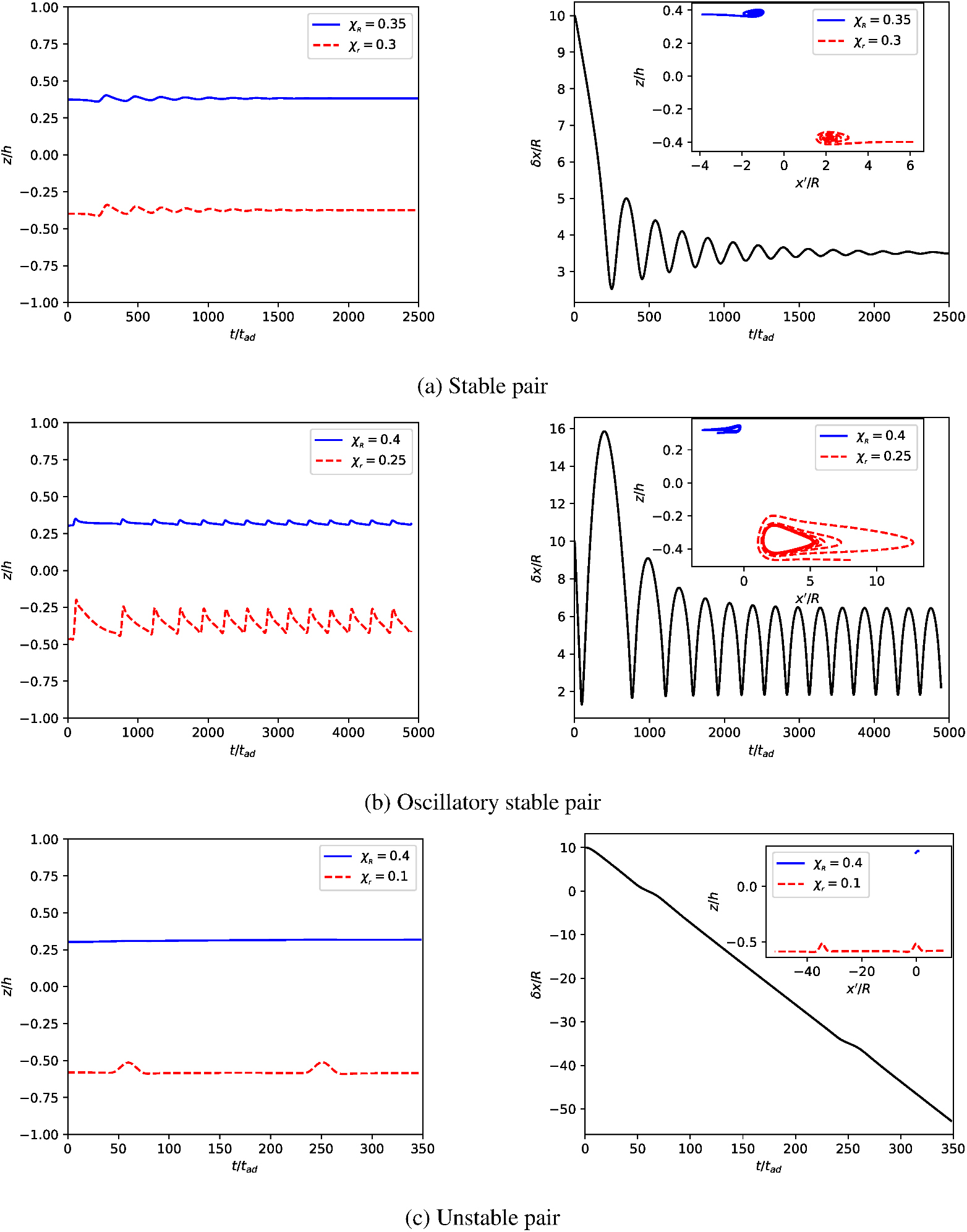
Types of particle pairs observed in this work. All examples are in staggered arrangement with the smaller particle initially leading and *δx*_0_/*R* = 10. The left column shows the lateral position of both particles as function of time. The right column displays the axial distance between both particles as function of time; the insets show the particle trajectories in the centre-of-mass system. The coordinate *x^′^* indicates the flow-wise coordinate with respect to the instantaneous centre of mass of the particle pair. (a) Stable pair of particles with *χ_R_* = 0.35 and *χ_r_* = 0.3. The inset shows that particle trajectories converge to a stable configuration. (b) Oscillatory stable pair with *χ_r_* = 0.4 and *χ_r_* = 0.25. The limit cycle of the trajectories is clearly visible in the inset. (c) Unstable pair with *χ_R_* = 0.4 and *χ_r_* = 0.1. The axial distance grows without bounds.

In contrast to a stable pair, an oscillatory stable pair is characterised by two particles whose axial distance *δx* keeps oscillating between two values within hydrodynamic interaction range. An example of a staggered oscillatory pair is shown in Fig. 2b for particles with *χ_R_* = 0.4 and *χ_r_* = 0.25. While the general dynamics is similar to that of the stable pair in Fig. 2a, the trajectories of the oscillatory stable pair converge to a limit cycle in the centre-of-mass frame.

An unstable pair is characterised by the axial distance *δx* increasing or decreasing without bounds. An example is shown in Fig. 2c for particles in the staggered arrangement with *χ_R_* = 0.4 and *χ_r_* = 0.1 where the smaller particle is initially leading. One reason for this instability is the large difference in axial particle speed due to a mismatch of the particles’ lateral equilibrium positions. The larger particle moves so much faster than the smaller particle that no stable pair can form during the time the particles interact hydrodynamically.

Since simulations are performed in a box that is periodic along the flow direction, pairs sometimes form between a particle and the periodic image of the other particle. A typical scenario is shown in Fig. 3 where the two particles are initially in a staggered arrangement (Fig. 3(a)). Under some circumstances, the leading particle moves away from the lagging particle and catches up with the periodic image of the other particle with which it consequently forms a pair (Fig. 3(b)). The actual axial distance between the initially considered particles is of the order *δx* ≈ L ≫ *R*, though. Initial conditions that lead to the formation of a periodic pair, rather than a stable or oscillatory stable pair as defined previously, are counted as initial conditions leading to an unstable pair since the initially considered particles move too far away from each other to form a pair. However, a periodic pair is still a stable (or oscillatory stable) pair that might have resulted from some unknown initial conditions under different circumstances. As such, periodic pairs are considered as (oscillatory) stable pairs as long as we are only interested in the existence of stable pairs and not in the circumstances under which they might have formed.

**FIG. 3:**
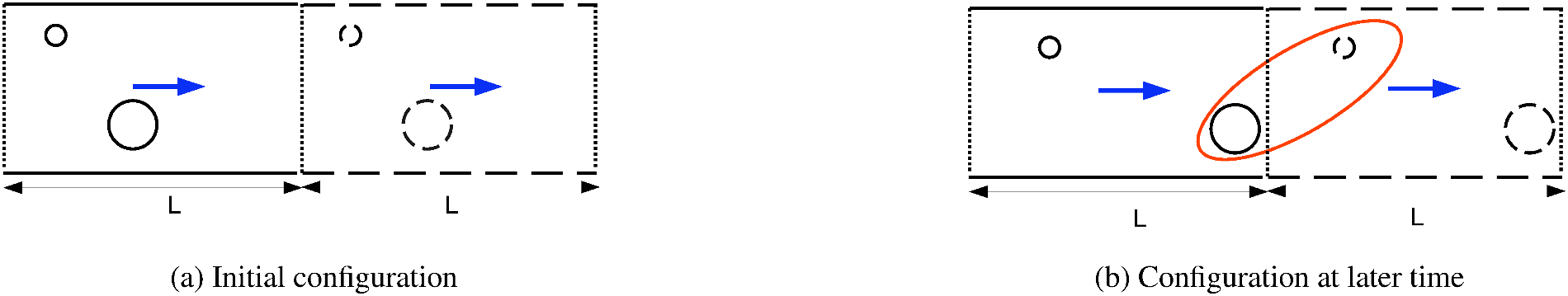
Schematic of the formation of a periodic pair. Solid lines represent the actual simulation domain and particles; dashed lines represent the adjacent downstream periodic unit cell. Dotted lines indicate the periodic boundaries. The flow is from left to right. (a) Initial configuration of two particles in the staggered arrangement. (b) Pair formation between one actual particle and the periodic image of the other particle at a later time.

From our initial simulations we know that particle pair formation depends on the particle sizes and the initial position of the particles. Thus, in the following, we study the effect of the confinement values (*χ_R_* and *χ_r_*) and the initial position of both particles on the formation and stability of particle pairs. We perform our analysis in three steps: (i) Assuming the absence of perturbations, particles are initially located far from each other and on their respective lateral equilibrium positions before they interact (Sec. IIIB). (ii) Pairs are initialised by putting the smaller particle in the vortex of the larger particle, and the stability of the pair is investigated (Sec. III C). (iii) Assuming that particles are generally affected by other particles in the channel, we investigate the effect of a perturbation of the initial conditions on the formation of pairs (Sec. III D).

### B. Pair formation from initially unperturbed particle positions

We start by assuming that both particles are initially far away from each other and had time to reach their individual lateral equilibrium positions, unperturbed by the possible presence of other particles in the channel. Since both particles have different sizes and, therefore, different lateral equilibrium positions, one of them is usually faster and therefore eventually catches up with the other particle.

Our initial simulations show that the particle pair is always unstable when the larger particle is initialised far downstream of the smaller particle since the larger particle has a lateral equilibrium position closer to the centreline and is, therefore, moving faster than the smaller particle. Thus, in all simulations in this section, we initialise the smaller particle as the leading particle. It is possible to form a stable pair with the larger particle leading if the initial lateral positions are different; we will investigate these cases in Sec. III D.

#### 1. Effect of initial axial distance

In order to ensure that particles are initially far away and not yet interacting hydrodynamically, we need to identify an appropriate initial axial distance, *δx*_0_. We consider two cases for a number of different values of *δx*_0_: (i) *χ_R_* = 0.3, *χ_r_* = 0.2 and (ii) *χ_R_* = 0.4, *χ_r_* = 0.2. The time evolution of the axial distance *δx* is shown in Fig. 4. For both cases, it can be seen that particles behave identically at late times as long as *δx*_0_ ≥ 8*R*. This behaviour is demonstrated more clearly in the insets of both panels where the time axis has been shifted in such a way that the first minimum of *δx*(*t*) occurs at *t’* = 0 for all curves with *δx*_0_ ≥ 8*R*. We conclude that *δx*_0_ = 8*R* ensures that particles do not interact initially. Hereafter, we initialise all simulations in this section with *δx*_0_ = 10*R*.

**FIG. 4:**
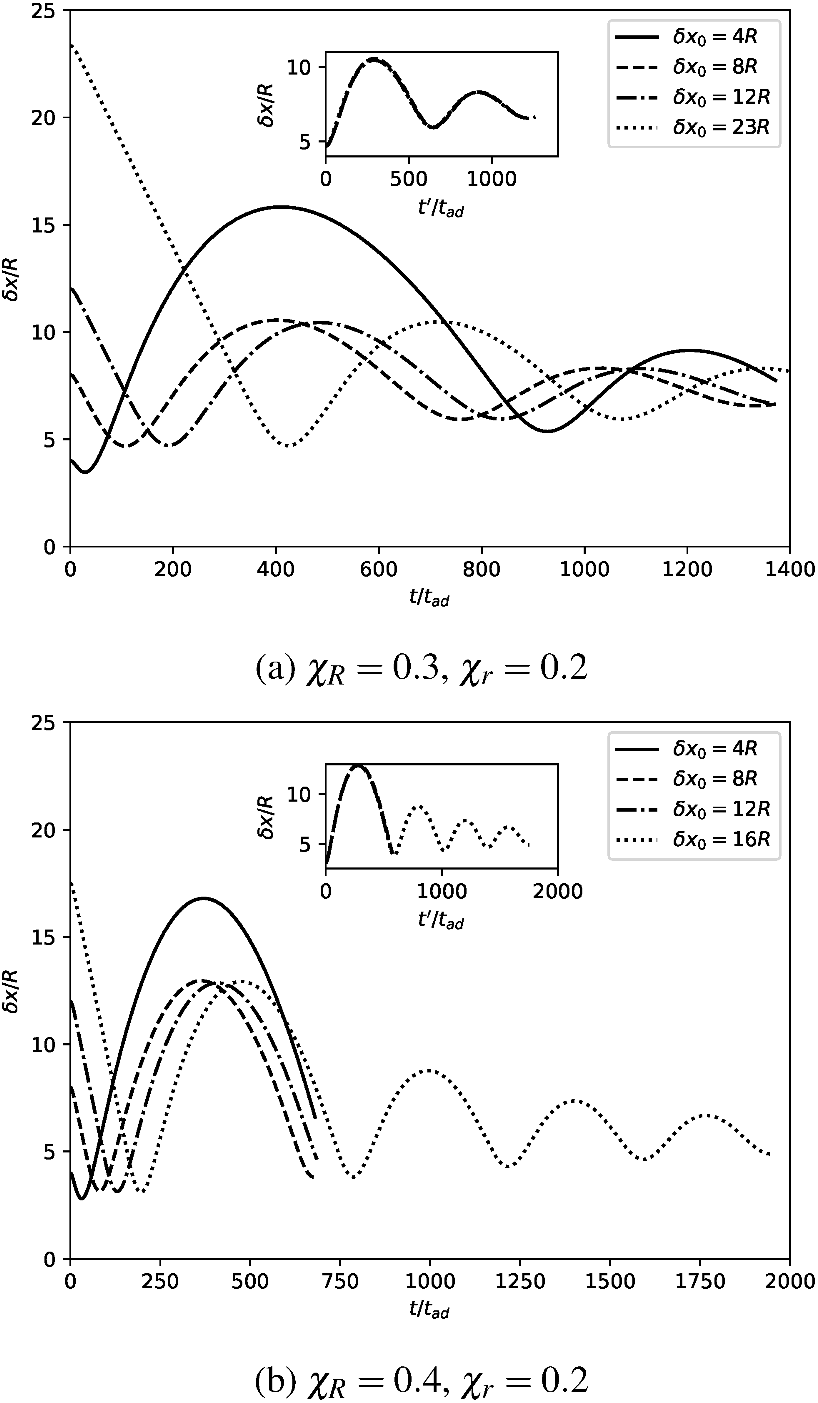
Time evolution of the axial distance between the particles for (a) *χ_R_* = 0.3, *χ_r_* = 0.2 and (b) *χ_R_* = 0.4, *χ_r_* = 0.2 for various initial axial distances *δx*_0_. The particle pairs behave identically at later times for *δx*_0_ ≥ 8*R*. The insets show the distance for *δx*_0_ ≥ 8*R* as function of shifted time *t’* such that the first minimum of *δx*(*t*) occurs at *t’* = 0.

#### 2. Effect of particle sizes on pair formation

Next we study the effect of particle size, *χ_R_* and *χ_r_* and size ratio *β* on the formation of stable pairs. Particles are initialised at their respective single-particle equilibrium positions.

The simulation outcomes are shown in Fig. 5. The green and grey shaded regions represent the smaller particle leading and lagging, respectively. We observe stable pairs (solid circles), oscillatory stable pairs (open circles) and no pair formation (crosses). As stated earlier, there is no pair formation when the smaller particle is initially lagging, both for staggered and linear arrangements. Since larger particles have equilibrium positions closer to the centre, these particles are in regions of faster flow and quickly move away from any lagging smaller particle. Since the initial distance between both particles is large, there is no hydrodynamic interactions between them, and the smaller particle has no way of forming a pair with the larger one. From now on, we only discuss those cases where the smaller particle is initially leading (green region).

**FIG. 5:**
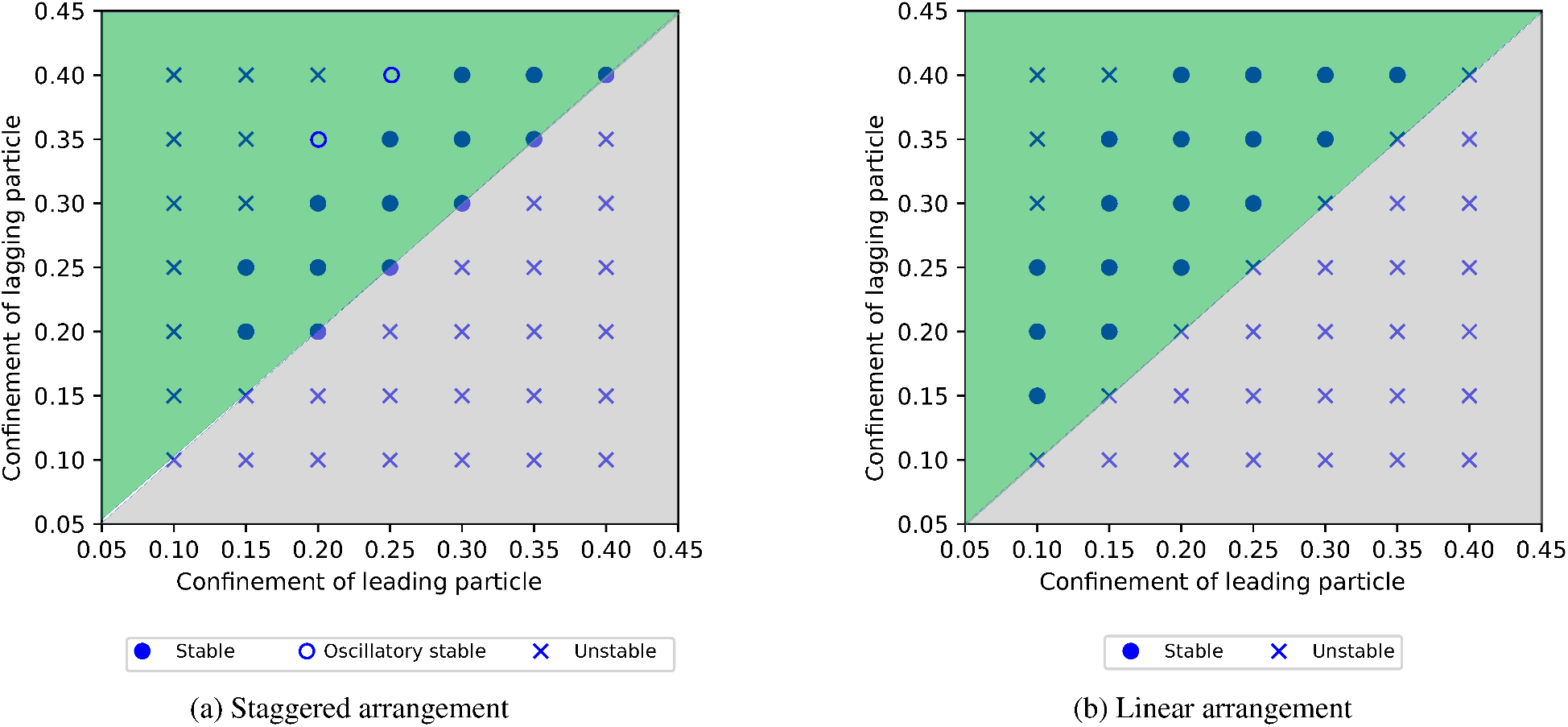
Effect of particle confinement on pair formation for particles initially far away from each other and at their respective lateral equilibrium positions. Particles are initialised either in the (a) staggered arrangement or (b) linear arrangement. The green and grey regions represent the smaller particle initially leading and lagging, respectively. Simulations are classified as one of three possible outcomes: stable pair (solid circle), oscillatory stable pair (open circle) and no pair (cross).

In the staggered arrangement (Fig. 5(a)), both particle size ratio and absolute particle size influence the outcome. When both particles are small compared to the channel, no pair formation is observed; the minimum confinement for which a pair forms is around *χ* = 0.15. Our streamline data (not shown here) suggest that the flow distortions caused by sufficiently small particles are not able to interact across the channel centreline, thus a stable staggered pair is unable to form when both particles are below a critical size. Furthermore, the particle size ratio *β* is an important factor: the particles do not form a pair when *β* ≈ 2 or larger. Importantly, the pair stability turns from stable to oscillatory stable before becoming unstable with increasing *β*. This observation suggests that oscillatory stable pairs can be found at the stability limit in terms of size ratio.

In the linear arrangement (Fig. 5(b)), pair formation predominantly depends on the particle size ratio *β*; there does not seem to be a minimum confinement for which pairs can form. In the limiting case of two identical particles (*β* = 1), we found that the axial distance between the particles increases steadily and apparently without upper bound. Since the increase in axial distance becomes progressively slower with distance, we aborted the simulation after *δx* reached about 15*R* without indication of reaching an equilibrium distance. This observation is in agreement with published results^20^. However, size heterogeneity enables the formation of linear pairs over a wide range of particle size ratios. Overall, we observe more stable pairs in the linear arrangement than in the staggered arrangement. Linear pairs can also form at larger particle size ratios, up to *β* ≈ 2.5.

To explore the role of size heterogeneity on pair formation close to the homogeneous limit, *β* ≈ 1, in the linear arrangement we have run simulations with varying degrees of mild size heterogeneity. Fig. 6 shows that a slight heterogeneity in particle size results in stable pairs forming. This observation is crucial since, in experiments, particle or cell properties are never perfectly homogeneous. The size ratio *β* also plays an important role for the equilibrium axial distance *δx*_eq_ in the pair as shown in the inset of Fig. 6. We observe a logarithmic divergence of *δx*_eq_ for *β* → 1. As a result, a small change in size heterogeneity for *β* ≈ 1 can lead to large variations in the axial distance. This observation explains the findings of Gao *et al*.^25^ who experimentally found that a small difference in particle size improves the particle focussing performance.

**FIG. 6:**
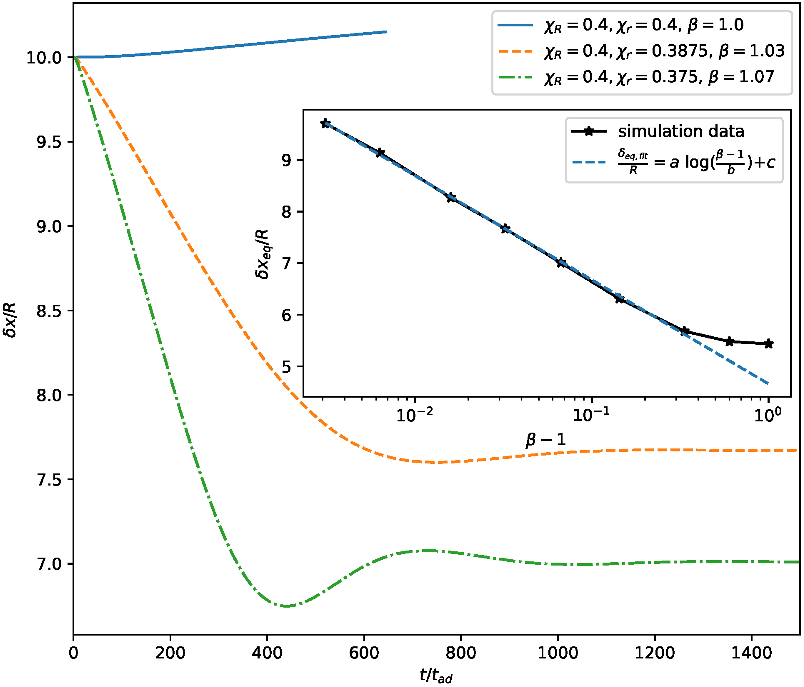
Linear pair formation for slightly heterogeneous particles. The time evolution of the axial distance *δx* between the particles shows that there is no pair formation for homogeneous particles (*β* = 1). However, a slight heterogeneity results in the formation of a pair. Variation of the equilibrium axial distance *δx*_eq_ with size ratio *β* is depicted in the inset; a logarithmic divergence is observed for *β* → 1 (fit parameters: *a* = −0.8756, *b* = 1.160, *c* = 4.531).

The equilibrium axial distance *δx*_eq_ is one of the most important properties of a particle pair. Fig. 7 shows the normalised axial distance for all observed stable pairs, both staggered and linear, as a function of the confinement of the smaller particle. Most notably, *δx*_eq_ is approximately twice as large for linear than for staggered pairs, which has been found previously^37^. Importantly, the equilibrium distance also depends on the individual particle sizes. Ignoring the dependence on *χ_R_*, staggered pairs tend to decrease their axial distance with increasing *χ_r_*, while we observe the opposite trend for linear pairs. For a given fixed value of *χ_R_*, however, staggered pairs show a weak increase of *δx*_eq_ with *χ_r_*. Furthermore, for a given fixed value of *χ_r_*, *δx*_eq_ of staggered pairs increases for decreasing *χ_R_*. For staggered pairs, there is no sign of a divergence of *δx*_eq_ for *β* → 1 in Fig. 7, which is in line with the observed stability of staggered homogeneous pairs.

**FIG. 7:**
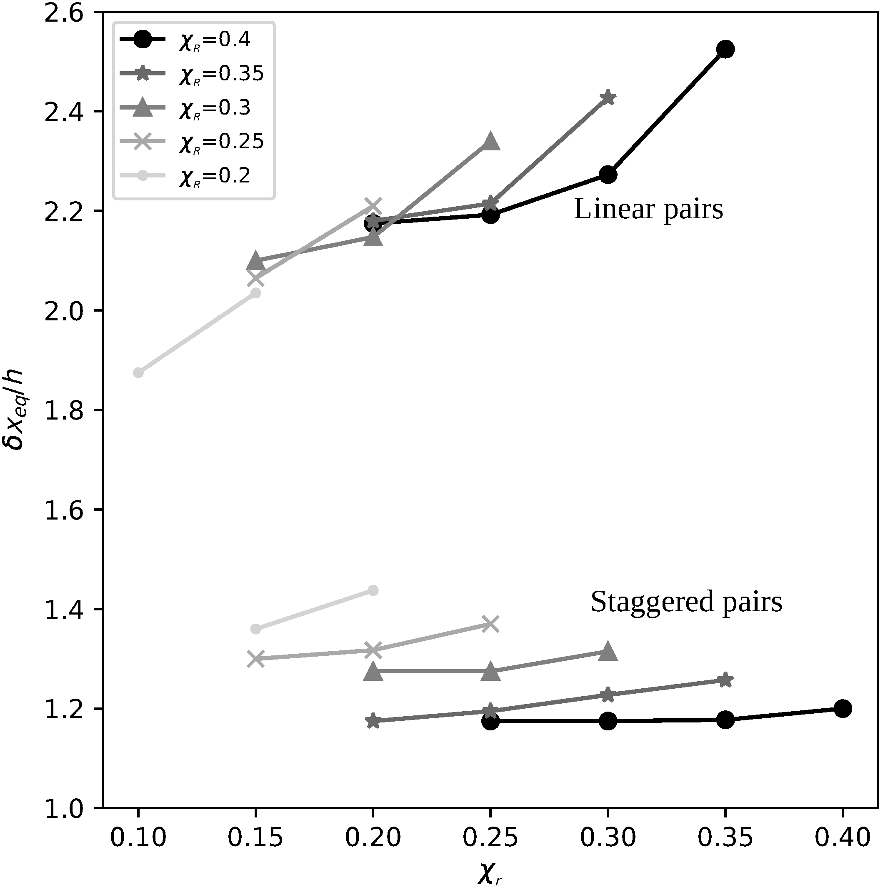
Normalised equilibrium axial distance *δx*_eq_/*h* for all stable pairs observed in Sec. IIIB as function of the confinement of the smaller particle, *χ_r_*. Note that, by definition, *χ_R_* ≥ *χ_r_* in all cases.

Fig. 8 shows the fluid streamlines around a single larger particle at its lateral equilibrium position in the co-moving frame of the particle. We observe that, in all our simulations, the smaller particle in a stable staggered pair eventually reaches the leading vortex caused by the larger particle. This effect has been described before^16,37^. For the linear stable pair, the smaller particle reaches the leading inner edge of the recirculation zone. We will take advantage of these observations in Sec. III C where we investigate the stability of already existing pairs.

**FIG. 8:**
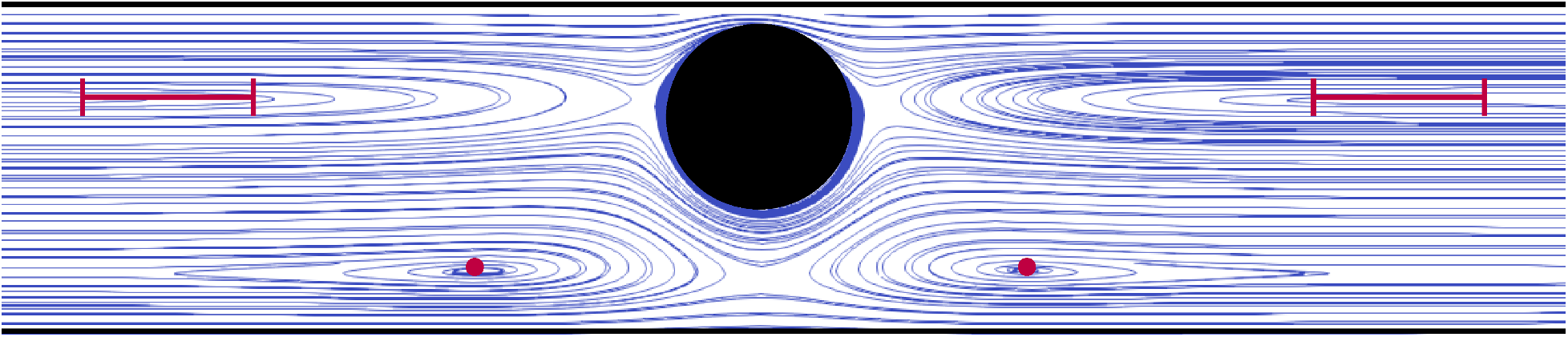
Streamlines (blue) in the co-moving frame of a single particle (black) at its lateral equilibrium position. Flow is from left to right. Red dots indicate the vortices located on the opposite side of the channel; a smaller particle reaches the leading vortex in case a stable staggered pair forms. The red bars highlight the inner edges of the recirculation zones on the same side of the channel; a smaller particle reaches the leading inner edge in case a stable linear pair forms.

Apart from the stable and oscillatory stable pairs defined in Fig. 2, we have also observed the formation of stable pairs involving periodic images. These periodic pairs were introduced in Sec. III A and play a special role since periodic images do not exist in the real world. However, the observation that stable periodic pairs can form for combinations of *χ_R_* and *χ_r_* that are marked as unstable in Fig. 5 implies that some pairs are able to form under different initial conditions than those assumed in this section. For example, we found stable periodic pairs in the linear arrangement with the smaller particle leading for *χ_R_* = 0.4, *χ_r_* = 0.15; *χ_R_* = 0.4, *χ_r_* = 0.1; *χ_R_* = 0.35, *χ_r_* = 0.1; and *χ_R_* = 0.3. *χ_r_* = 0.1 which are all marked unstable in Fig. 5. Based on the observation of stable pairs that do not form when particles are initially at their lateral equilibrium positions and far away from each other, we need to investigate which particle arrangements are stable (assuming that pairs already exist) and under which initial conditions additional stable pairs can form in the first place. We will address both questions in Sec. III C and Sec. III D, respectively.

### C. Stability of already existing pairs

The initial conditions used in Sec. III B are very specific, and we have already seen that not all possible stable pairs are formed from these initial conditions. Thus, in this section, we study the stability of pairs already existing. To this end, we initialise particles in such a way that the larger particle is at its lateral equilibrium position and the smaller particle is located either in the vortex of the larger particle (staggered arrangement) or at the edge of the recirculation zone of the larger particle (linear arrangement), see Fig 8.

There are two distinct cases for the linear and staggered arrangements, respectively: the smaller particle can initially be leading or lagging since there are two vortices and two recirculation zones. Once the pair is initialised, we track the axial distance *δx* to determine whether the pair is stable or not. The resulting stability map for both staggered and linear pairs is shown in Fig. 9.

**FIG. 9:**
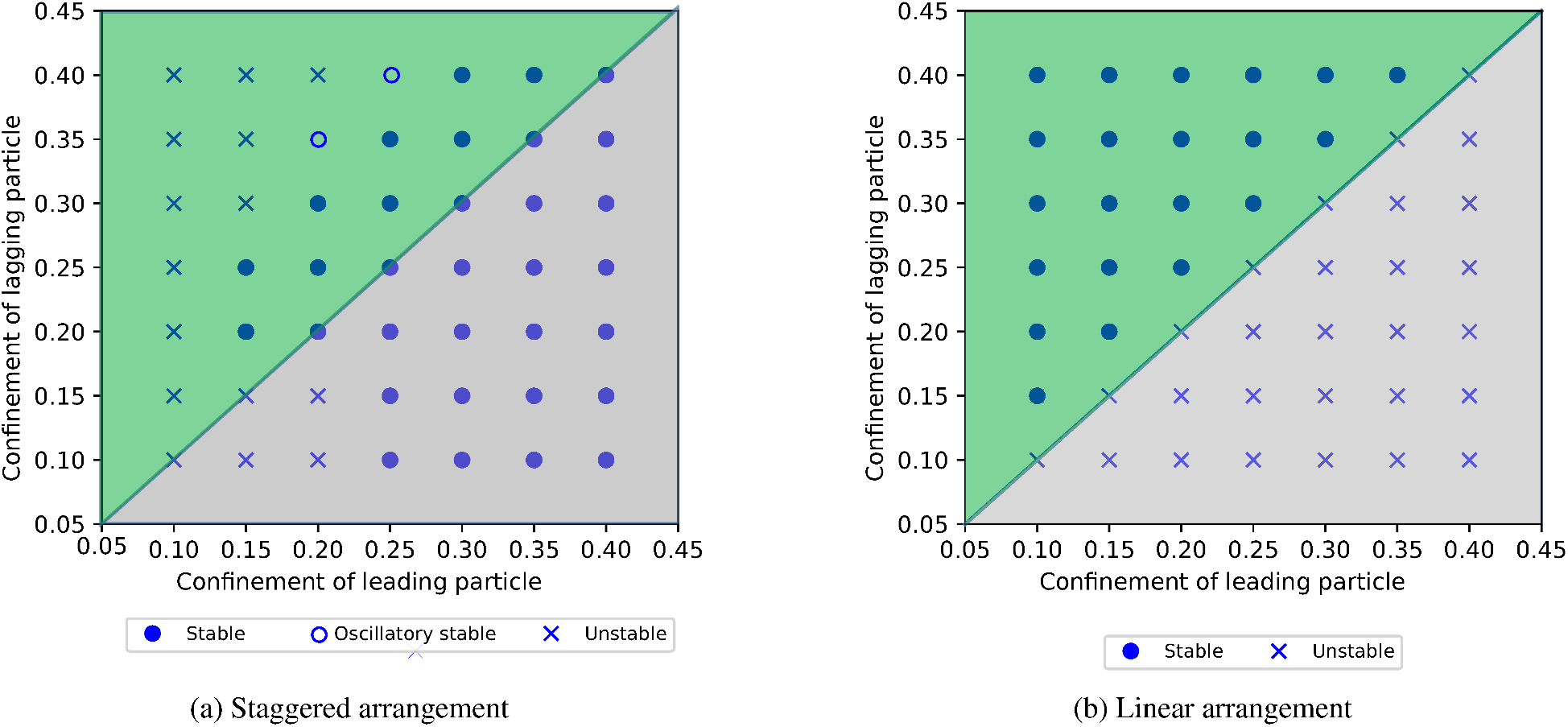
Effect of particle confinement on stability of pairs initialised according to Fig. 8 in the (a) staggered arrangement and (b) linear arrangement. The green and grey regions represent the smaller particle initially leading and lagging, respectively. Simulations are classified as one of three possible outcomes: stable pair (solid circle), oscillatory stable pair (open circle) and no pair (cross).

For the cases where the smaller particle is leading in the staggered arrangement (green area in Fig. 9a), we did not find any more stable pairs than those already observed in Fig. 5a. However, there exist stable pairs when the larger particle is leading in the staggered arrangement in Fig. 9a, despite Fig. 5a not showing any of these stable pairs. These new stable pairs are not very robust and require careful selection of the initial positions to be observed. Although these pairs are stable according to our definition, they are sensitive to perturbations that offer the leading larger particle the opportunity to move away due to its preferred lateral equilibrium position closer to the centreline where the flow is faster.

In the linear arrangement, particle pairs are stable when the smaller particle is leading and unstable when the larger particle is leading for all investigated combinations of size ratio (Fig. 9b). These findings are similar to those in Fig. 5b, although pairs with a large value of *β* = *R/r* did not form when particles were initially far away from each other. These results are in contrast to the findings of Chen *et al*.^26^ who reported that linear pair formation is only possible when the larger particle is leading. However, their simulations were in 2D, omitting transverse flow effects which Haddadi and Morris^42^ demonstrated alter the stability characteristics within the pair^42^.

We have seen that the initial configuration of the particles plays an important role. Some pairs tend to be more robust and can form when particles at their respective lateral equilibrium position approach each other for the first time, while other stable pairs require more carefully chosen initial conditions. Since the initial conditions in our simulations reflect an arbitrary instant in time in a real-world application where particles might be in arbitrary locations, we need to understand better how the choice of initial conditions affects pair formation.

### D. Pair formation from initially perturbed particle positions

We showed in Sec. III B that pairs of particles can form for specific combinations of particle sizes when both particles are initialised at their respective lateral equilibrium positions. We then discovered in Sec. III C that more particle pairs, if already formed, are stable. Based on these findings, we can distinguish between three possible scenarios. For a given combination of confinement values *χ_R_* and *χ_r_* (and therefore *β*), exactly one of the following outcomes is true:

1. A pair forms when particles are initially at their respective lateral equilibrium positions. This scenario is typically expected when the suspension is very dilute and particles have the time to migrate to their lateral equilibrium positions before encountering each other. These are combinations of particles that are marked as stable in Fig. 5.
2. A pair forms when particles are initially at other lateral positions. This situation is expected to be more common in less dilute suspensions where particles are perturbed by the presence of other particles such that particles cannot migrate to their lateral equilibrium positions before they encounter each other. These are combinations of particles that are marked as stable in Fig. 9 but as unstable in Fig. 5.
3. No pair forms independently of the initial particle positions. These are combinations of particles that are marked as unstable in Fig. 9.

In this section, we explore the second scenario. It would be unfeasible to scan the entire space of initial positions of both particles, thus we pursue a different strategy. Since larger particles are less perturbed by the presence of other particles and migrate faster toward their lateral equilibrium position^43^, we assume that the larger particle is initially at its lateral equilibrium position. The smaller particle, however, is released at different lateral positions to mimic the effect of an upstream perturbation. Furthermore, both particles are still initialised on the mid-plane (*y* = 0) to reduce the complexity of the problem, and *δx*_0_ = 10*R* in all cases.

We investigate a single combination of particles with the smaller particle lagging in the staggered arrangement that was found stable (Fig. 9) but did not result in a pair when initialised at the lateral equilibrium positions (Fig. 5): *χ_R_* = 0.4, *χ_r_* = 0.1 (*β* = 4). The trajectories of both particles in the centre-of-mass system are shown in Fig. 10 for a range of initial positions of the smaller particle. The dashed line corresponds to the case where the smaller particle is initially at its lateral equilibrium position, and the square and the thick dots represent the positions of the particles in the stable arrangement. It is particularly obvious that the smaller particle, when it is initially at its lateral equilibrium position, is far away from its stable point in the pair and immediately moves away from the larger particle. This observation confirms the finding of Patel and Stark^23^ who reported that the presence of the second particle can change the stability of the single-particle equilibrium positions. Thus, for a stable pair to form, we hypothesise that the smaller particle should be close to the lateral position of its stable point when both particles approach each other. Indeed, there exists a ‘stability band’ of initial lateral positions (indicated by the red line) that all result in the formation of the same stable pair. All investigated cases of initial positions outwith the stability band do not result in the formation of a stable pair. Since the width of the stability band is small compared to the height of the channel, we expect that the probability of this specific pair forming is relatively low in a real-world scenario where the smaller particle could be at a random location initially. This stability band could explain the formation of heterogeneous pairs after a number of passing interactions in a simulation using periodic boundary conditions reported by Li *et al*.^24^. On each pass, the lateral position of the lagging particle is modified due to the previous passing interaction. We hypothesise that, for the instance the pair actually forms, the lagging particle approaches the leading particle within the stability band. In the future, it could be investigated how the width of the stability band is related to the probability of pair formation.

**FIG. 10:**
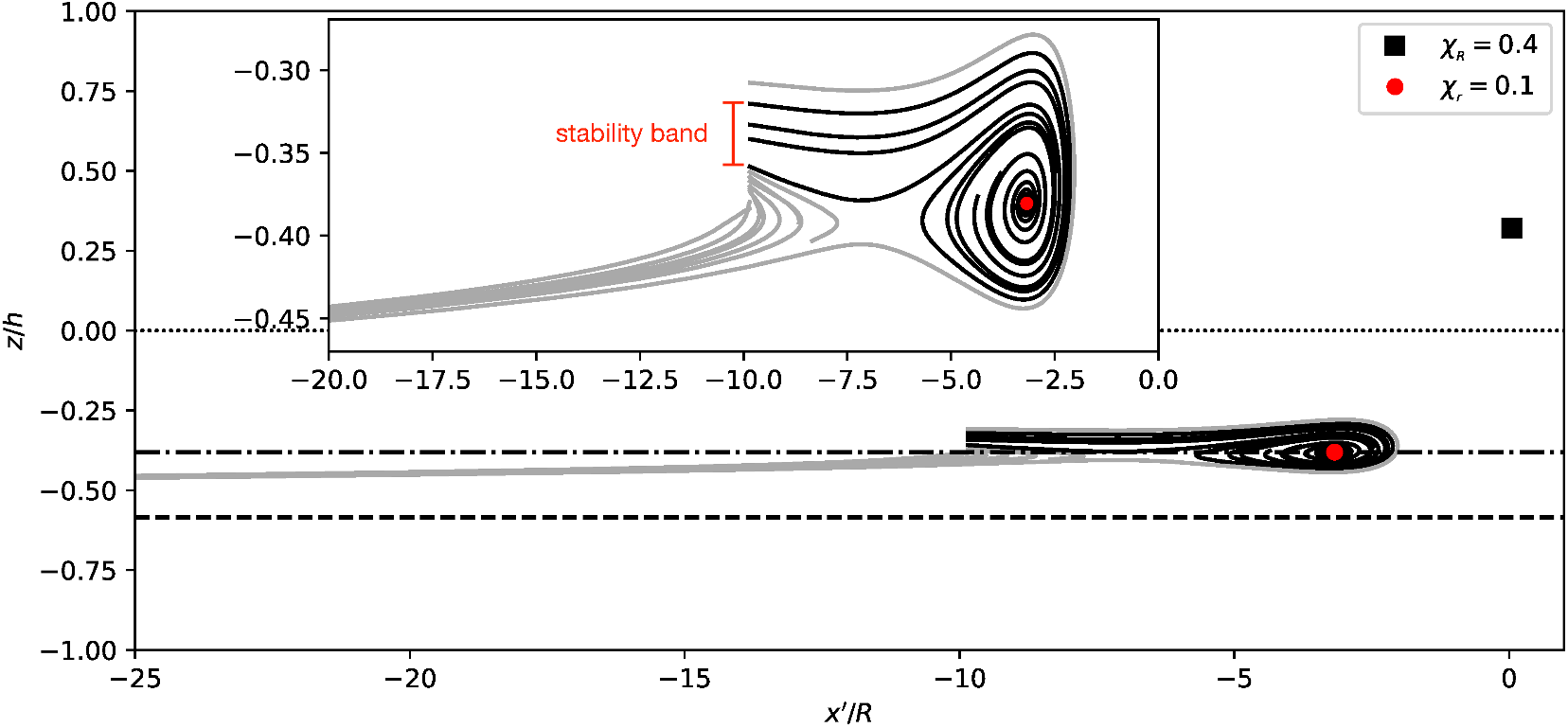
Particle trajectories are shown in the centre-of-mass system for *χ_R_* = 0.4 and *χ_r_* = 0.1 where the smaller particle is lagging in the staggered arrangement. Flow is from left to right. Square and red dot represent the particle positions in the stable pair with the larger particle on the right near *x’* = 0. The dash-dotted line indicates the lateral position of the smaller particle in the stable pair. Particles are initialised with *δx*_0_ = 10*R*, thus trajectories of the smaller particle start near *x’* = −10*R*. The dashed line shows the lateral equilibrium position of the smaller particle, leading to no pair formation. The inset shows the zoomed-in region around the stable position of the smaller particle. The dotted line represents the centerline. A stable pair only forms when the smaller particle is initialised in the ‘stability band’ denoted by the red measurement line.

The example case considered here shows that the initial conditions play an important role in the formation of a pair of particles with different sizes. While a considerable number of pairs can form when particles are initially at their respective lateral equilibrium positions, other pairs can only form under different initial conditions. The presence of other particles in a channel might lead to the perturbations needed to enable the formation of pairs that would not form under unperturbed conditions.

## IV. SUMMARY AND CONCLUSIONS

The formation of stable particle pairs is crucial for various inertial microfluidic applications, such as particle focussing and separation. Identifying the conditions under which stable particle pairs form is important for designing microfluidic devices. Despite the use of inertial microfluidics to separate cells and particles of different sizes, there is a paucity of studies investigating the formation of pairs of particles of different sizes. The present work addresses this need by investigating the effect of the size and initial position of particles on the formation and stability of heterogeneous pairs under moderate inertia.

We used an in-house immersed-boundary-lattice-Boltzmann-finite-element solver to simulate a pair of particles of different sizes (radii *r* and *R* ≥ *r*) and identical softness (Laplace number 100) moving in a pressure-driven flow (along the *x*-axis) through a rectangular channel (half-width w and half-height *h* < *w*) at Reynolds number 10. Particles were initially located on the plane defined by the mid-points of the longer edges (*y* = 0) along the channel width. While particles move along the channel, they generally undergo lateral migration along the height axis (*z*-axis) and all particle motion occurs on the *x*-*z*-plane midway between the side walls. We considered two different particle arrangements: staggered (particles on opposite sides of the channel centreline) and linear (particles on the same side of the channel centreline). To model particles in a dilute suspension that approach each other from a large distance, particles were initialised with an axial distance *δx*_0_ = 10*R* for which hydrodynamic interactions are weak.

First, we considered pair formation for configurations where both particles are initially at their respective lateral equilibrium positions. These configurations are denoted ‘unperturbed’ since they would be expected when particles are isolated and have enough time to migrate laterally before encountering each other. We found that stable pairs in the staggered and linear arrangements form for a wide range of confinement values *χ_R_*=*R/h* and *χ_r_* = *r/h* and their ratio *β* = *R/r*. The pair formation strongly depends on which particle is leading and lagging. No pair formation was observed when the smaller particle is lagging since the larger particle is generally located closer to the centreline where it moves faster. When the smaller particle is leading, staggered pairs only form when both particles have a minimum size while linear pairs can form even when both particles are small compared to the channel height. Furthermore, pairs only form if the size ratio *β* is sufficiently small (typically *β* < 2–3); this critical size ratio is larger for linear pairs than for staggered pairs. We also confirmed earlier studies showing that no stable linear pairs form when both particles have the same size (*β* = 1). However, even a mild size heterogeneity of the order of a few percent leads to stable linear pairs with a large axial spacing δx_eq_. This finding is important since particles or cells are never perfectly uniform in experiments. Generally, the axial spacing *δx*_eq_ in stable pairs is roughly twice as large in linear pairs than in staggered pairs, and *δx*_eq_ depends on both particle sizes.

Second, we investigated the stability of already existing pairs, independently of their possible formation, by placing the smaller particle in one of the eddies caused by the larger particle. While all pairs observed forming in the first part of our study were confirmed to be stable, we also identified additional stable pairs that were unable to form under unperturbed conditions. In particular, staggered pairs are stable over a wide range of particle sizes even when the leading particle is larger than the lagging particle. However, we did not find any stable linear pairs when the leading particle is larger than the lagging particle.

Third, in order to understand why not all possible stable pairs form when particles are initialised at their respective lateral equilibrium position, we performed another study to investigate the effect of the initial position of the smaller particle on pair formation. We found that there exists a ‘stability band’: a finite range of initial lateral positions of the smaller particle that lead to stable pairs (as long as the pair is stable for the given combination of particle sizes). This stability band might or might not include the lateral equilibrium position of the smaller particle, thus explaining why some pairs are inaccessible under unperturbed initial conditions. Our findings imply that upstream perturbations caused by the presence of additional particles, as would be expected even in dilute suspensions in inertial microfluidics, might play an important role in the formation of stable pairs that would not be able to form for two unperturbed particles.

Our study was performed with slightly deformable particles in straight channels at Reynolds number 10. It remains an open question how the mechanisms of pair formation are different in curved channels at higher Reynolds number as they are often used in inertial microfluidic applications. We hope that this study creates additional impetus for the simulation-informed design of inertial microfluidic devices for particle focussing and separation.

## AUTHOR DECLARATIONS

### Author Contributions

**Krishnaveni Thota:** conceptualization (equal), methodology (equal), formal analysis (lead), visualization (lead), writing — original draft (lead), writing — review and editing (equal). **Benjamin Owen:** conceptualization (equal), methodology (equal), formal analysis (supporting), visualization (supporting), writing — original draft (supporting), writing — review and editing (equal), project administration (supporting). **Timm Krüger:** conceptualization (equal), methodology (equal), formal analysis (supporting), visualization (supporting), funding acquisition, writing — original draft (supporting), writing — review and editing (equal), project administration (lead).

### Data availability

The data that support the findings of this study are available from the corresponding author upon reasonable request

## Acknowledgement

This work used the Cirrus UK National Tier-2 HPC Service at EPCC (http://www.cirrus.ac.uk).

For the purpose of open access, the authors have applied a Creative Commons Attribution (CC BY) licence to any author accepted manuscript version arising from this submission.

